# Viral and bacterial traits associated with the success of the yellow pencil coral, *Madracis mirabilis,* in Curaçao’s coral reefs

**DOI:** 10.1101/2025.04.04.647251

**Authors:** Bailey A. Wallace, Natascha S. Varona, Alexandra K. Stiffler, Mark J.A. Vermeij, Cynthia Silveira

## Abstract

Coral reefs have experienced extensive coral loss and shifts in community composition worldwide. Despite this, some coral species appear naturally more resistant, such as *Madracis mirabilis* (herein *Madracis*). *Madracis* has emerged as the dominant hard coral in Curaçao, comprising 26% of coral cover in reefs that declined by 78% between 1973 and 2015. Although life history traits and competitive mechanisms contribute to *Madracis*’ success, these factors alone may not fully explain it, as other species with similar traits have not shown comparable success. Here, we investigated the potential role of microbial communities in the success of *Madracis* on Curaçao reefs by leveraging a low-bias bacterial and viral enrichment method for metagenomic sequencing of coral samples, resulting in 77 unique bacterial metagenome-assembled genomes and 2,820 viral genomic sequences. Our analyses showed that *Madracis*-associated bacterial and viral communities are 1.24-fold and 1.61-fold richer than the communities of five sympatric coral species combined. The *Madracis*-associated bacterial community was dominated by *Ruegeria* and *Sphingomonas*, genera that have previously been associated with coral health, defense against pathogens, and bioremediation. The viral community exhibited a 50% enrichment of proviruses relative to the viral communities of other corals. These proviruses have the genomic capacity to laterally transfer genes involved in antibiotic resistance, central metabolism, and oxidative stress responses, potentially enhancing the adaptive capacity of the *Madracis* microbiome and contributing to *Madracis*’ success on Curaçao’s reefs.

**IMPORTANCE:** Understanding why some coral species persist and thrive while most are in fast decline is critical. *Madracis mirabilis* is increasingly dominant on degraded reefs in Curaçao, yet the role of microbial communities in its success remains underexplored. This study highlights the potential role of *Madracis-*associated bacterial and viral communities in supporting coral resilience and competitive success. By identifying key microbial partners and viral genes that may enhance host stress tolerance and defense against pathogens, we broaden the understanding of how the coral holobiont contributes to species persistence under environmental stress. These insights are valuable for predicting reef community shifts in a changing climate and open avenues for microbiome-informed strategies to support coral conservation and restoration.

## INTRODUCTION

A promising approach to coral reef conservation and restoration is the identification and exploration of coral species and reefs that are doing better than expected, given the pressures they are under (1). These interesting cases may provide insights into the physiological, genetic, and microbial adaptations that enable survival and stability under environmental stressors like warming, acidification, or reduced water quality. In the Caribbean, *Madracis mirabilis*^1^ (commonly known as yellow pencil coral) exemplifies this resilience among dramatic losses in coral cover throughout the region. The coral reefs of Curaçao are no exception when it comes to the global trends of reef decline over the past few decades (2). Anthropogenic stressors have contributed to reduced coral cover, elevated algal presence, and shifts in community composition (3). Despite a 78% decline in hard coral coverage from 1973 to 2015 in Curaçao, *Madracis mirabilis* (*Madracis*) has become the dominant hard coral species, accounting for 26% of coral cover and even increasing in some shallow reef sites (4–6).

Across Curaçao’s reef, *Madracis* experiences the highest growth and percent cover in areas proximal to chronic anthropogenic stressors, including nutrient loading, pollution, and organic matter inputs (6, 7). Typically, these stressors lead to community-level shifts and increasingly microbialized reefs, which are detrimental to coral health (8, 9). Microbialization is stimulated by organic matter inputs resulting in the proliferation of copiotrophic bacteria, which contribute to the formation of hypoxic zones through elevated heterotrophic respiration and often include opportunistic coral pathogens (9). Despite these challenges, *Madracis* demonstrates remarkable resilience, showing relative insensitivity to coral diseases that affect many other species. Life history traits such as fast growth rates and high population turnover, make weedy species of coral, like *Madracis*, particularly resilient to the impacts of habitat degradation (10). As an efficient heterotroph, *Madracis* consumes zooplankton, bacteria, and particulate matter from the water column, which may offset the typical decreases in autotrophic feeding (i.e., photosynthesis) caused by environmental stressors (11–13). *Madracis* not only thrives under environmental and microbial stress but also coexists with a dense community of benthic competitors. Its unique morphology, where tissue at the base of each branch recedes as the colony expands, creates space for diverse cryptic communities of crustose coralline algae, sponges, algal turfs, cyanobacterial mats, and other benthic organisms. This proximity to competitors may explain its heightened aggression through mesenterial filaments and sweeper tentacles (7). While these life history, nutrition, and aggression traits likely contribute substantially to *Madracis*’ expansion and dominance in Curaçao, numerous coral species share similar characteristics without achieving comparable success. Thus, these traits may not fully explain *Madracis*’ disproportionate success.

Microorganisms of the coral holobiont are both crucial to coral fitness and highly sensitive to the physiological status of their host (14). The composition of microbial communities within the coral holobiont is shaped by the dynamic interactions between these symbiotic microorganisms and environmental conditions, where the most beneficial community, given the environmental context, is selected (15). Dysbiosis in coral-associated microbial communities has been identified as a key contributor to disease (16, 17) with resident bacteria acting directly as opportunistic pathogens under certain environmental pressures (18–22). Other microbiome members provide significant benefits to their coral hosts, as highlighted by the Beneficial Microorganisms for Corals (BMCs) concept (23, 24). Beneficial microbes are thought to outcompete harmful microbes and even reverse dysbiosis within the coral holobiont through key functions like pathogen control, neutralization of toxic compounds, and nutrient cycling (25–29). Though microbial community composition can differ widely across coral species, flexibility in microbial partnerships may provide advantages to corals on short timescales, in contrast to genetic mutation and natural selection (15). The ability of microbes to act both as pathogens and as agents of adaptability highlights the complex dynamics of coral-microbe relationships and their potential to determine competitive success.

Viruses may also play a significant role in modulating microbial dynamics within corals. Bacteriophages, viruses that infect bacteria, act as agents of horizontal gene transfer and exert top-down control on bacterial communities (30–32). In other systems, viral predation can determine the colonization success of invading bacterial strains (33) and function as a form of mucosal immunity (32, 34, 35). Viruses can also carry auxiliary genes that can modulate host metabolisms, virulence, and other phenotypes (31, 36). Long-term relationships between viruses and bacteria through viral genome integration (forming a prophage) can provide the bacterium with several advantages, such as immunity to superinfection by related phages and the acquisition of novel genes, which can spread beneficial traits like antibiotic resistance or virulence factors (31). Moreover, prophage persistence can help maintain genetic diversity within bacterial populations, promoting evolutionary flexibility and adaptation in fluctuating environments (37, 38).

Here, we describe the bacterial and viral communities associated with *Madracis mirabilis*, the corals it interacts with, and seawater from the coral boundary layer to uncover the potential roles of these microbes in *Madracis*’ ecological success. We hypothesize that the *Madracis* microbiome displays unique traits related to community diversity and genetic makeup that can contribute to supporting this coral’s persistence.

## MATERIALS AND METHODS

### Sample collection

Images, coral biopsies, and coral boundary layer (CBL) seawater samples were collected at 12 sites along the southwestern coast of Curaçao in June of 2022 (Fig. S1A; Table S1). At the time of sampling, *Madracis* was the dominant scleractinian coral across surveyed reef sites on Curaçao, visibly dominating the benthic reefscape (Fig. 1A, Fig. 1B). *Madracis* colonies were frequently observed in direct competition with other benthic organisms, including neighboring coral species (Fig. 1C, Fig. 1D). At each site, a *Madracis* patch interacting with one of five other coral species was sampled, as shown in Figure S1B. Interactions were imaged at high resolution for quantification of interaction zone outcomes (N = 11 with one site lost; see supplementary methods for details). Coral biopsies of approximately 1 cm^3^ (containing mucus, tissue, and skeleton) were collected using a chisel and hammer and placed in Ziplock polyethylene bags with ambient seawater. Samples were placed on ice during transfer to the laboratory at the CARMABI research station (maximum 1h transfer time), where ambient seawater was removed, and coral samples were flash-frozen and stored at -80°C until later processing. For comparison with the microbiome in the surrounding environment, CBL seawater samples were collected from four areas around each *Madracis* patch using a 500 ml syringe equipped with a 30 cm-long flexible tube: within the *Madracis* fingers (Inter-branch; IB1), the boundary layer overlaying the *Madracis* (within 10 cm of the coral surface; BL1), the boundary layer over the interaction zone (BL2) and the boundary layer of an upstream coral of the same interacting species (BL3) (Fig. S1B).

**Figure 1.**
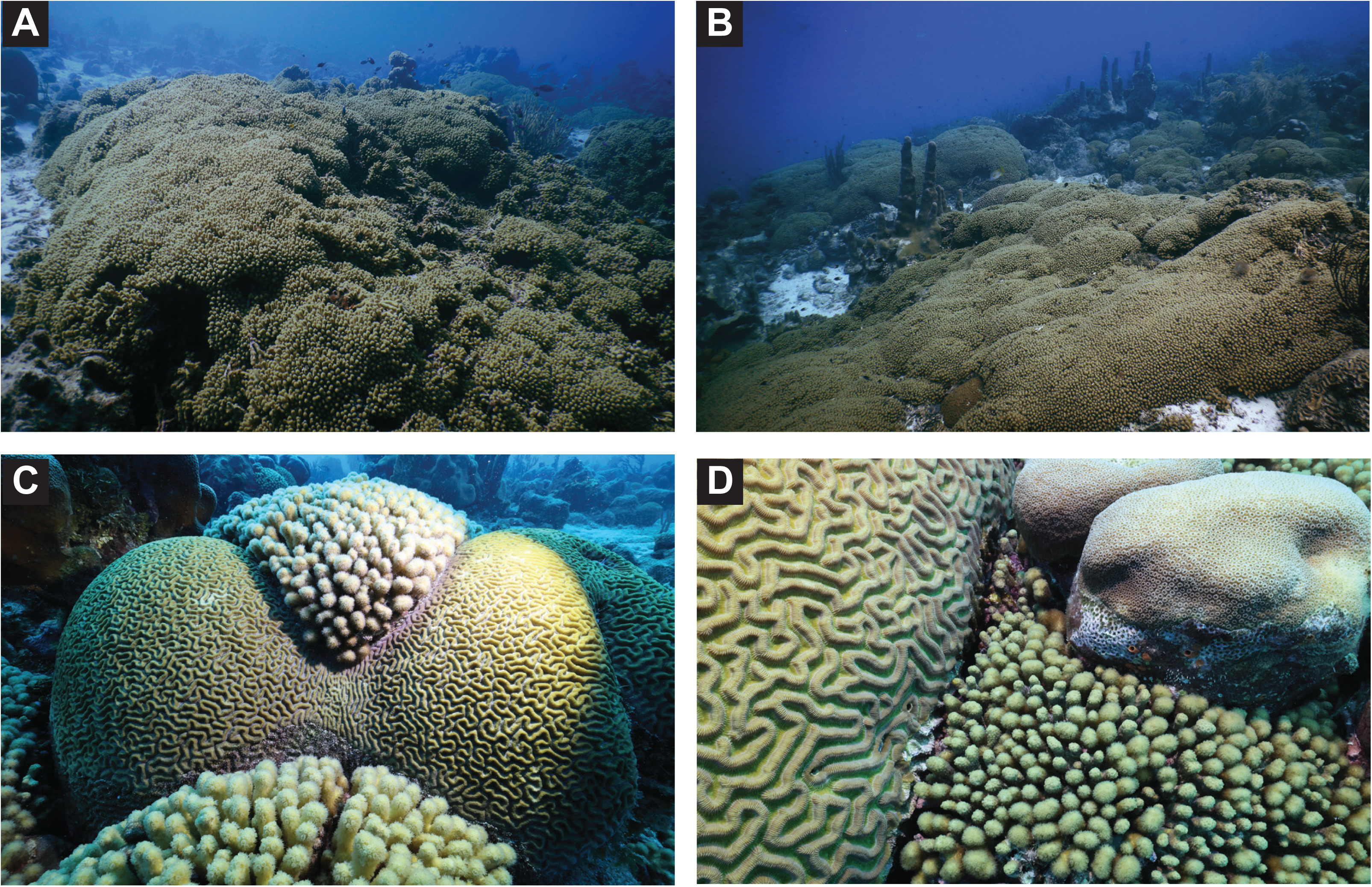
*Madracis* colonies in the reefscape and their interactions with other benthic organisms. (A, B) *Madracis* colonies widely distributed across the reefscape, contributing significantly to coral cover. (C, D) *Madracis* colonies interacting with neighboring coral species, highlighting competition for space.

### Epifluorescence microscopy of coral boundary layer seawater

For epifluorescence microscopy, 1 mL of raw CBL sample was fixed with paraformaldehyde (2% final concentration; Thermo Scientific Chemicals, Waltham, MA, USA) and vacuum-filtered onto a 0.02 µm Anodisc (Cytiva, Marlborough, MA, USA). Anodiscs were air-dried and stored flat at -20°C during transportation to the University of Miami, FL. In the laboratory, nucleic acid-containing particles on the filters were stained by placing the filter on a 100 µl drop of SYBR Gold Nucleic Acid Gel Stain (10X final concentration; Invitrogen, Waltham, MA, USA) for ten minutes. Excess stain was removed by wetting the Anodisc from below with two drops of 100 µL molecular grade water and gently blotting the bottom of the filter with a lint-free wipe. Filters were air-dried for 30 minutes without exposure to light and mounted between a glass microscope slide and coverslip with 20 µL of mounting solution (ascorbic acid (0.1%), phosphate-buffered saline (1X), and 0.02 µm-filtered glycerol (50%)). Slides were visualized using oil immersion at 630 X magnification on a ZEISS Axio Imager.A2 using ZEN image processing software (ZEISS, Oberkochen, BW, GER). For each slide, 10 images for BL1, BL2, and BL3 and five images for IB1 were captured as technical replicates using an Axiocam 506 mono camera. The mean number of cells and virus-like particles (VLPs) across these fields of view was scaled to the total area of the Anodisc, yielding the count of cells and VLPs per mL in each CBL sample. Two sites, “Kokomo Beach” and “Tugboat Beach” were removed from the analysis due to low-quality slides, which affected their reliability for downstream analyses.

### DNA extractions and sequencing

For the CBL seawater metagenomes, four 500 mL samples from each site were 8.0 µm-filtered (polycarbonate track-etched membrane filter; Whatman, Buckinghamshire, UK) and precipitated overnight with 10% Polyethylene Glycol 8000 (PEG; Fisher BioReagents, Pittsburgh, PA, USA). The next morning, PEG-precipitated bacteria and viruses were collected on a 0.22 µm Sterivex filter (MilliporeSigma, Burlington, MA, USA) and frozen at –20 °C for later processing. DNA from Sterivex filters containing CBL seawater samples from 6 sites (N = 21) was extracted using a modified Nucleospin Tissue Kit protocol (Macherey-Nagel, Düren Nordrhein-Westfalen, Germany) (39). Briefly, the Sterivex filters were thawed and air-dried by forcing air through with a syringe. A pre-lysis step was performed by capping one end of the filter and adding 360 µL of Buffer T1 and 50 µL of proteinase K directly to the Sterivex, followed by overnight incubation at 56°C on a rotating spit. The next day, 400 µL of Buffer B3 was added, and the samples were incubated at 70°C for 30 minutes. Lysates were extracted from the Sterivex filters using a luer lock syringe and transferred to 2 mL tubes. The procedure continued from step 4 of the NucleoSpin Tissue kit protocol for human or animal tissues and cultured cells. DNA libraries were prepared using the Nextera XT DNA Library Preparation Kit (Illumina) and IDT Unique Dual Indexes using 1 ng of DNA. Fragmentation was performed with Illumina Nextera XT fragmentation enzyme, followed by PCR amplification (12 cycles) and purification with AMpure magnetic Beads (Beckman Coulter). Libraries were quantified using a Qubit 4 fluorometer (ThermoFisher Scientific, Waltham, MA, USA) and a Qubit dsDNA HS Assay Kit (Invitrogen, Waltham, MA, USA). Libraries were sequenced on a NovaSeq platform 2x150bp (Illumina, San Diego, CA, United States).

DNA extractions from 24 coral samples followed a host depletion protocol based on tissue disruption, DNAse treatment, and size fractionation (40, 41) to enrich viral and bacterial communities. Genomic DNA was quantified using the Qubit 2.0 Fluorometer (ThermoFisher Scientific, Waltham, MA, USA) and libraries were prepared with the NEBNext Ultra™ II DNA Library Prep Kit for Illumina following the manufacturer’s recommendations. Genomic DNA was fragmented by acoustic shearing (Covaris S220), cleaned, end-repaired, and adapter-ligated, followed by PCR enrichment. Libraries were validated using a High Sensitivity D1000 ScreenTape on the Agilent TapeStation (Agilent Technologies, Palo Alto, CA, USA) and quantified by real-time PCR (Applied Biosystems, Carlsbad, CA, USA). Libraries were sequenced on an Illumina NovaSeq X Plus instrument 2x150bp. *Orbicella faveolata* samples (N = 19) collected in Curaçao in 2021 and extracted and sequenced with the same method were also incorporated into this dataset, expanding the total number of sampling sites by three (Fig. S1A). The collection and sequencing details of the 2021 samples were previously reported (41).

### Identification of bacterial and viral sequences

Raw metagenomic reads were adapter-trimmed and quality-filtered (trimq=30, maq=30) with BBDuk (v39.01) (42), generating over 900M quality-controlled, paired-end reads as quantified by FastQC (Andrews, S., 2010). For bacterial community analysis, metagenomic reads were analyzed for taxonomic classification using Kaiju v1.10.1 with the proGenomes Database v3 (43, 44). The relative abundances of bacterial genera were calculated using the following equation:

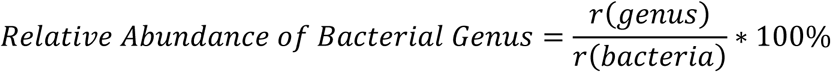

where *r(genus)* is the total number of reads matched to a particular bacterial genus, and *r(bacteria)* is the total number of reads matched to the domain Bacteria. Bacterial metagenome-assembled genomes (bMAGs) were generated by combining and improving the single-sample coverage binning outputs from MaxBin2 v2.2.7 (45), MetaBat2 v2.15 (46), and CONCOCT v1.1.0 (47) with metaWRAP v1.2.1(48). Bins with ≥50% completion and ≤10% contamination (N = 2,112) were classified with GTDB-Tk v2.4.0 (GTDB release 220) (49). The anvi_genome_similarity and anvi-dereplicate-genomes modules of Anvi’o v8 (50) were used to dereplicate bMAGs at a similarity threshold of 95%, generating 77 species-level representative bMAGs. To identify proviruses within these genomes, VIBRANT v1.2.1 (51) and geNomad v1.7.4 (52) were run separately on the bMAGs. A phylogenetic tree of the representative bMAGs was constructed using the GTDB-Tk infer module. For viral community analysis, quality-controlled reads from each sample were assembled with metaSPAdes v3.15.5 using default parameters (53). The resulting assemblies were analyzed using the ViWrap v1.3.1 pipeline (54), utilizing both VIBRANT and geNomad viral identification tools. Within this pipeline, viral contigs were binned using VRhyme v1.1.0 (55) and taxonomically classified using the NCBI RefSeq viral protein, VOG HMM, and IMG/VR v4 databases (56–58). All resulting binned and unbinned viral genomes and genome fragments identified with the two distinct viral identification pipelines were pooled and dereplicated at MIUViG standards (95% ANI over 80% AF) with CheckV’s (v1.0.1) rapid genome clustering based on pairwise ANI (59), resulting in 13,113 unique viral sequences. These viruses were further quality-filtered by removing all sequences with no viral genes and CheckV quality “Not-determined”, resulting in a final database of 2,820 viruses. Reads from each sample were mapped at 95% identity to the de-replicated viral and bMAG databases with SMALT v0.7.6, and relative fractional abundances of viral sequences and bMAGs were calculated using the following equation:

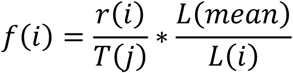

Here, *r(i)* is the number of reads mapped to each viral or bacterial genome, *T(j)* is the total number of reads mapped in a given sample, *L(mean)* is the mean genome length in the dataset, and *L(i)* is the length of the mapped sequence (60). Only sequences with >10 reads mapped were considered present in a sample. Genome annotation of viruses and bMAGs was performed with MetaCerberus v1.4.0 (61). The most abundant functional gene categories and genes encoded by proviruses and bMAGs were identified by calculating their average relative abundance across coral groups. Functional gene abundances were derived from the abundances of the genomes encoding each function, normalized within each sample, and averaged across the two coral groups (*Madracis* and others). For bMAGs, MetaCerberus gene products were filtered using keywords to identify genes previously identified as relevant to bacteria-host symbioses. Best hits were further filtered by database to select the 15 most differentially abundant gene pathways, COG (all available pathways) and KEGG (filtered for pathways containing the keyword “metabolism”), between *Madracis* and other corals. *Madracis* indicator viruses (described below) encoding integrases (N = 3) were visualized with genoPlotR (62). In addition, two high-quality genomes of *Caudoviricetes* proviruses identified by CheckV were selected for genome visualization.

### Statistical analysis

Differences between groups in the microscopy-based abundance data were tested with an analysis of variance (ANOVA), followed by Tukey HSD tests to identify significant pairwise differences. To assess the similarity between viral and microbial community profiles, the Bray-Curtis similarity algorithm was applied to bacterial genus relative abundances and viral relative fractional abundances (vegan: vegdist) and visualized with non-metric multidimensional scaling (vegan: metaMDS). Significant differences between groups were identified using Adonis permutational ANOVAs (adonis2; 999 permutations) and pairwise Adonis tests (pmartinezarbizu: pairwise.adonis2). The homogeneity of multivariate dispersion was evaluated on the Bray-Curtis distance matrices (vegan: betadisper). Significant differences in NMDS ordination of bacterial and viral communities were observed when grouped by sample type (coral vs. CBL seawater) and year (2021 vs. 2022), yet these differences may also be due to the coral species (Fig. S2; Table S2; Table S3; Table S4).

To describe which bacterial genera and viruses contributed to the between-group differences identified by the PERMANOVA tests, we used similarity percentages (vegan: SIMPER) analysis. Only the 20 bacterial genera and viral sequences with the highest contribution to the dissimilarity between coral groups were reported. To identify which viral community members had high specificity and fidelity to the *Madracis* community, indicator species analyses (ISA) were used (indicspecies: multipatt). Diversity metrics, including Shannon diversity, Simpson diversity, Evenness, and Richness, were calculated on the bacterial genera and viral genomes in each group (vegan: diversity). To test the effects of seven coral samples that yielded significantly fewer sequencing reads (< 1M reads) and *Orbicella faveollata* (OFAV) samples collected in a different year (with different beta-dispersion in bacterial [p = 0.0011, Tukey HSD] and viral [p = 0.0005, Tukey HSD] communities), we removed those samples and reanalyzed the dataset, finding similar results (Table S5, Table S6; Table S7; Table S8).

### Data availability

Coral and CBL seawater metagenomic sequence data for this project are available on NCBI (PRJNA1214393 and PRJNA1061506). The code used to conduct this analysis is publicly available on GitHub (https://github.com/Silveira-Lab/Wallace_Madracis_Holobiont). R code and data are available on figshare (doi:10.6084/m9.figshare.28216004), along with assembled viral genomes (doi:10.6084/m9.figshare.28259492), and QC reports (doi:10.6084/m9.figshare.28215692).

## RESULTS

### Bacterial and viral communities in coral boundary layer seawater

Interbranch seawater from between the *Madracis* fingers (IB1; N = 10) had significantly higher viral abundance (ANOVA: F(3, 33) = 11.03, p = 3.59e-05), bacterial abundance (F(3, 33) = 20.99, p = 8.62e-08), and virus-to-microbe ratios (VMR; F(3, 33) = 8.65, p = 0.00023) compared to all boundary layer samples collected above the corals (BL1, N = 10; BL2, N = 7; and BL3, N = 10; Fig. 2; Table S9). Viral and bacterial abundances in IB1 were 26- to 31-fold and 10- to 12-fold higher, respectively, while VMRs were 3-fold higher than in BL samples. No significant differences were detected among BL1, BL2, and BL3 (Table S10). Despite differences in cellular and viral abundances between the interbranch and boundary layer samples, bacterial community composition only differed between the interbranch samples (IB1) and the boundary layer overlaying the *Madracis* (BL1; F(1, 9) = 3.42, p = 0.025; Table S11). Viral communities were also similar among boundary layer (BL) samples but differed significantly between IB1 and both BL1 (F(1, 9) = 4.27, p = 0.003) and BL2 (F(1, 8) = 1.87, p = 0.017). Combined boundary layer and interbranch communities were distinct from those associated with coral tissue (Table S3; Table S4). Given the similarity among boundary layer communities, we combined all seawater samples into a single coral boundary layer category, termed “CBL seawater”, for downstream comparisons.

**Figure 2.**
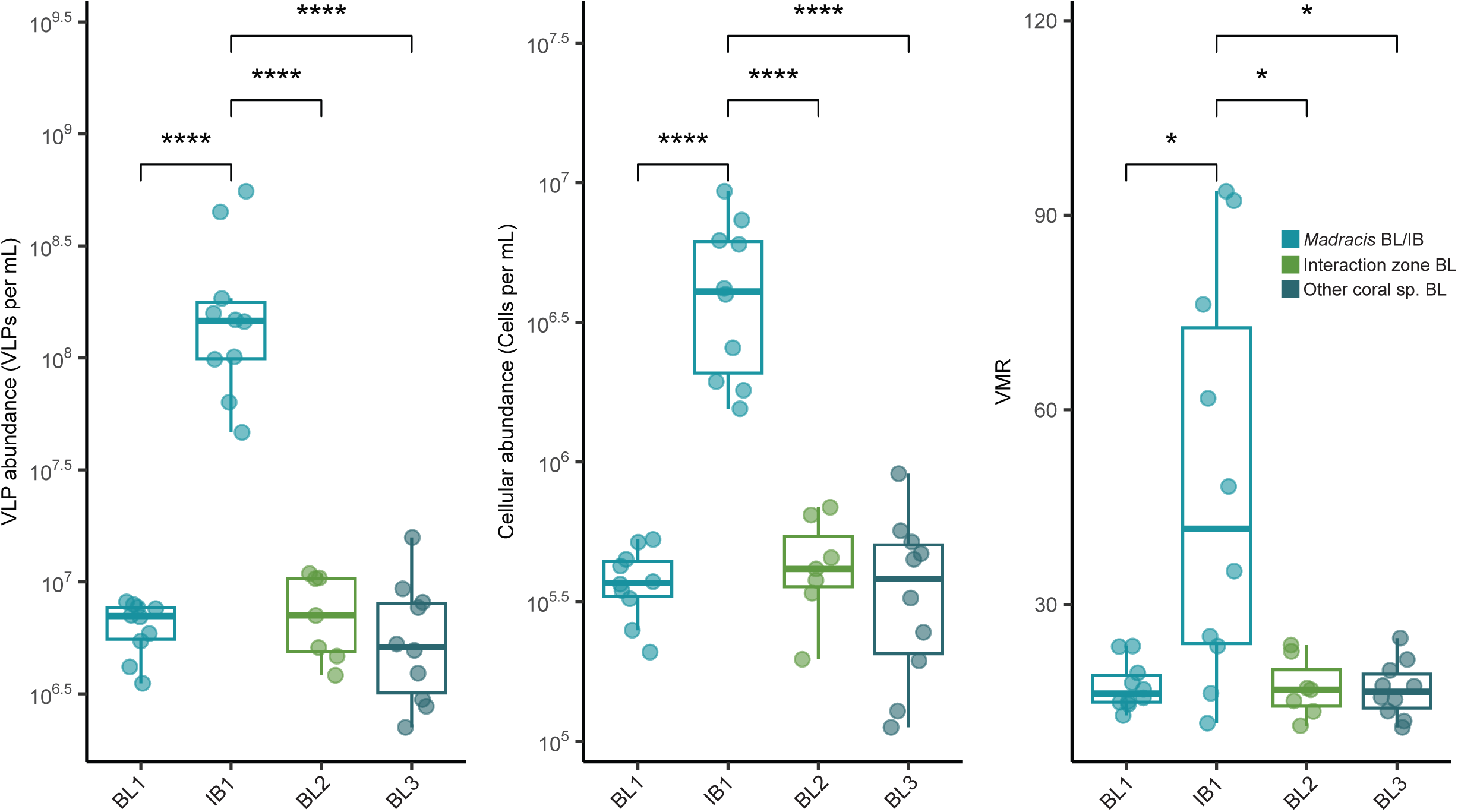
Microbial abundances in coral boundary layer and interbranch seawater. Epifluorescence microscopy counts of Viral-like particles (VLPs), Cells, and the virus-to-microbe ratio (VMR) for seawater samples spanning *Madracis* interactions. Mean count per mL of sample and VMRs are displayed for each of the four seawater sample types including the *Madracis* boundary layer (BL1), *Madracis* interbranch (IB1), interaction zone boundary layer (BL2), and other coral species boundary layer (BL3). Only statistically significant differences are displayed (level of significance is represented by asterisks).

### Interaction outcomes

To assess whether bacterial and viral communities influenced the outcomes of direct competition between *Madracis* and other coral species, we quantified these interactions according to Fig. S3A. *Madracis*’ interaction outcomes were generally not species-specific, but other coral species lost more against *Madracis* (Fig. S3B) than against other benthic substrates (Fig. S3C). Bacterial and viral communities were not significantly different in winning or losing interaction outcomes (F(1, 22) = 0.896, p = 0.511 for bacteria and F(1, 22) = 1.571, p = 0.139 for viruses; Fig. S4). Ratios of *Bacteroidetes* to *Bacillota* (previously known as *Firmicutes*), which are markers of dysbiosis (63–66), were similar between winning and losing corals (Fig. S5). Due to the lack of differences in competition outcomes, we did not pursue this line of inquiry in subsequent analyses.

### Bacterial communities associated with Madracis

Bacterial community assemblages differed among *Madracis*, other coral species, and the combined CBL seawater (F(2, 61) = 28.184, p = 0.001; Fig. 3A). Beta dispersion of the bacterial communities was 1.9-fold greater for *Madracis* and 1.8-fold greater for other coral species combined compared to the CBL seawater (p adj <0.0001 for both comparisons; Tukey HSD) but was similar between the bacterial communities of *Madracis* and other coral species (p adj = 0.91, Tukey HSD; Fig. S6). The mean Shannon diversity index of the CBL seawater bacterial community (2.85 ± 0.14) was ∼1.6-fold and ∼1.4-fold lower than that of *Madracis* (4.50 ± 0.25; p adj < 0.0001) and other corals (4.03 ± 0.08; p adj < 0.0001), respectively (Fig. 3B; Table S7). This pattern was similar for Simpson diversity, where the CBL seawater bacterial community (0.69 ± 0.03) was ∼1.4-fold lower than that of *Madracis* (0.94 ± 0.01; p adj < 0.0001) and ∼1.3-fold lower than that of other coral species (0.93 ± 0.01; p adj < 0.0001). The mean richness of bacterial genera was highest in the *Madracis* community (1053.25 ± 17.55), compared to that of all other coral species combined (852.94 ± 36.94; p adj = 0.02, Tukey HSD), but was not significantly higher than the bacterial richness in the CBL seawater (878.71 ± 55.49; p adj = 0.06, Tukey HSD). The evenness of the bacterial communities was similar between *Madracis* (0.65 ± 0.04) and other coral species (0.60 ± 0.01; p = 0.40, Tukey HSD) but was significantly lower in the CBL communities (0.043 ± 0.03; p < 0.0001 for both comparisons).

**Figure 3.**
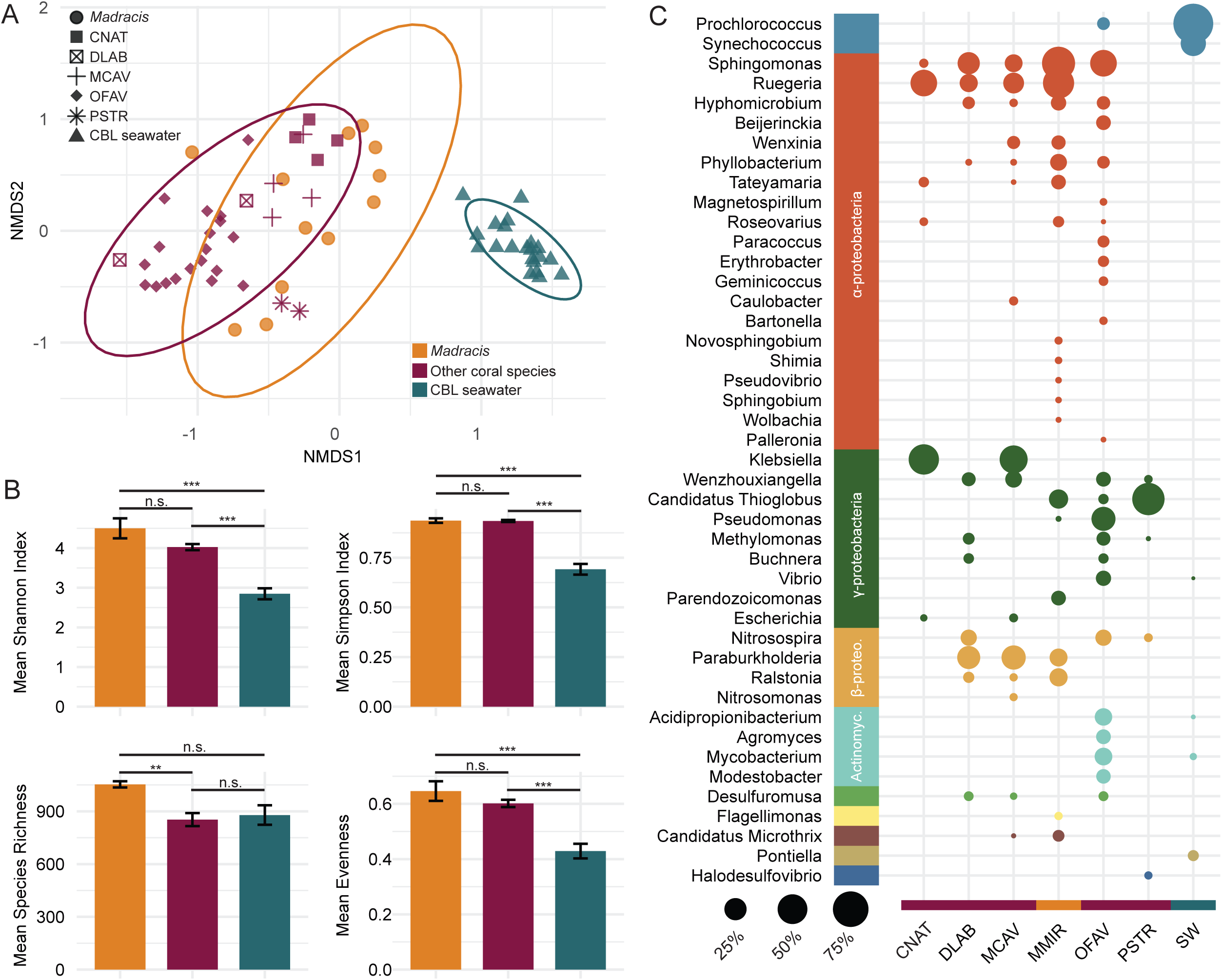
Bacterial community distinctions between *Madracis*, all other coral species in this study, and coral boundary layer (CBL) seawater. (A) Non-metric multidimensional scaling (NMDS) plot by bacterial genus. NMDS plot is based on Bray Curtis distances calculated from the relative abundance of reads and utilizes 999 permutations. Circles denote a 95% CI for the clustered points. (B) Mean diversity indices of bacterial communities for each of the three groups, including Shannon Index, Simpson Index, Evenness, and Richness of bacterial genera. Significance codes: 0 ‘***’, 0.001 ‘**’, and 0.01 ‘*.’ (C) Bubble plot displaying the relative abundances of bacterial genera by coral species/seawater for genera representing > 1% of the bacterial community. The size of the circles indicates percent relative abundance, and the color of the bubbles indicates bacterial classes. Coral species names are abbreviated as follows: *Colpophyllia natans* (CNAT), *Diploria labyrinthiformis* (DLAB), *Montastraea cavernosa* (MCAV), *Madracis aurentenra* (MAUR), *Orbicella faveolata* (OFAV), *Meandrina meandrites* (MMEA), and *Pseudodiploria strigosa* (PSTR).

A total of 1,099 bacterial genera across 61 classes were identified in the combined dataset through our read-based analysis. Of these, 44 genera represented at least 1% of bacterial relative abundance within a given sample and are reported as the mean relative abundance of each genus in each coral species or the CBL seawater (Fig. 3C). *Madracis* bacterial communities were dominated by *Ruegeria* (8.38% ± 2.39%; mean ± SE) and *Sphingomonas* (6.14% ± 2.27%). In contrast, other coral species were, on average, dominated by the genera *Klebsiella* (8.60% ± 1.80%), Candidatus *Thioglobus* (7.33% ± 3.58%), and *Pseudomonas* (7.13% ± 0.57%). The twenty genera making the largest contribution to the dissimilarity between the bacterial communities of *Madracis* and other coral species accounted for 27% of the total variation between groups (Table S12), with 13 having a higher abundance in *Madracis*, including *Ruegeria*, *Hyphomicrobium*, and *Wenxinia*. Seven genera had abundances at orders of magnitude lower levels in *Madracis*, including *Klebsiella*, *Nitrosospira*, *Wenzhouxiangella*, and *Beijerinckia* (Table S12).

A total of 2,112 refined bacterial metagenome-assembled genomes (bMAGs) were dereplicated at a 95% similarity, yielding 77 representative species-level bins with 83.90% ± 1.63% completion and 1.91% ± 0.21% contamination (Fig. S7). These 77 bMAGs spanned 14 phyla, 19 classes, and 49 genera of bacteria (Fig. 4; Table S13). 78% (N = 60) of the bMAGs were identified to the genus level by GTDB-Tk and 26% (N = 20) contained proviruses. The most abundant bMAG across all coral samples was *Endozoicomonas*_WF-CRL-4_bin.66, representing an average of 31.54% ± 7.02% of the bMAGs relative abundance in *Madracis* and 42.92% ± 5.35% in other corals. *Poriferisocius*_SM-CRL-3_bin.41 (8.87% ± 3.36% in *Madracis* and 10.31% ± 2.53% in other corals), *Sphingomonas*_SNAKE-CRL-3_bin.123 (5.42% ± 3.85% and 11.53% ± 2.91%), and a bin of the class *Bacteroidia* (Unknown_DR-CRL-4_bin.142; 4.49% ± 2.11% and 9.03% ± 2.60%) followed in abundance (Fig. 4).

**Figure 4.**
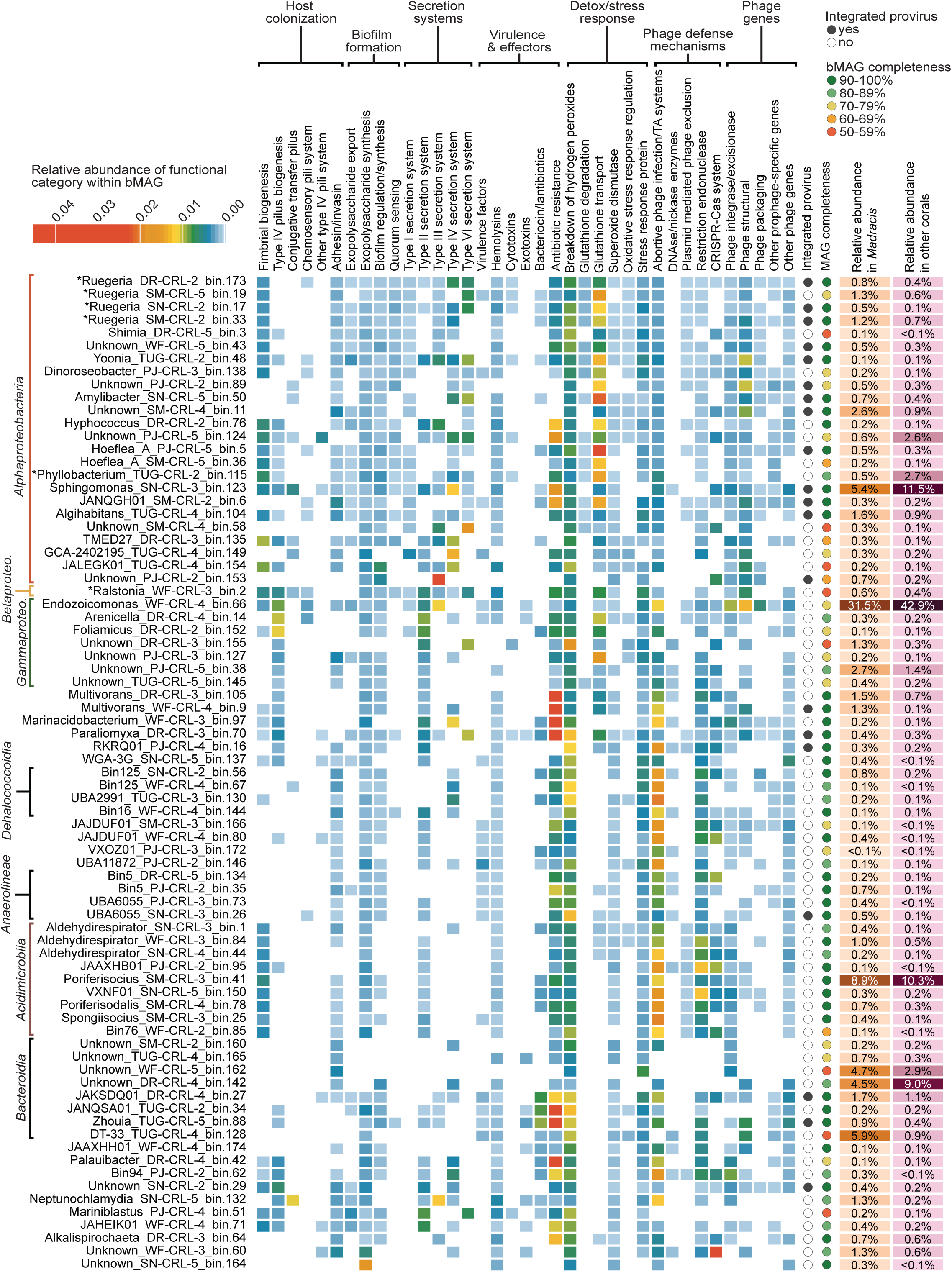
Symbiotic functions of bacterial metagenome-assembled genomes (MAGs) from corals. Functional analysis of the 77 representative bacterial metagenome-assembled genomes (bMAGs). The color gradient from light blue to red represents the relative abundance of each functional category within a given bMAG, as determined by the summed relative abundance of gene products within each functional category. Genomes are further annotated with the presence/absence of an integrated prophage (identified by VIBRANT and/or GeNomad), the completeness of the genome (ranging from 51.05% to 100%), and the relative abundance of each genome in *Madracis* and in other coral species. On the y-axis, genomes are ordered by their phylogenomic distance, as determined by GTBD-Tk. Classes with more than four representative genomes are described along the y-axis, and the full taxonomic classification of each bMAG can be found in Table S13.

Functional analysis of the 77 representative genomes revealed 178,108 individual gene hits, spanning 8,845 unique gene products (Fig. 4). Gene products involved in fimbrial and pilus biogenesis, biofilm formation, secretion systems, effector/toxin production, and detoxification/stress responses were found throughout the bMAGs. Additionally, numerous genes related to phage defense mechanisms, particularly toxin-antitoxin (TA) systems and restriction endonucleases, were identified in the majority of the representative genomes. Although confirmed prophages were detected in only 26% of bMAGs (N = 20), many more of these bacterial genomes encoded viral structural proteins (66%; N = 51) and phage integration genes (80%; N = 61), suggesting a potentially higher prevalence of prophages. Three genera distinguishing the bacterial communities of *Madracis* and other corals in the SIMPER analysis had representative bMAGs, including *Ruegeria* (N = 4), *Ralstonia* (N = 1), and *Phyllobacterium* (N =1). *Ruegeria* genomes were enriched in Type IV and VI secretion systems, antibiotic resistance mechanisms, genes related to the breakdown of hydrogen peroxide, and glutathione transport genes (Fig. 4). Three of the four *Ruegeria* genomes were lysogens. The *Ralstonia* genome contained similar genes, along with a broader diversity of pilus biosynthesis genes related to eukaryotic host colonization. *Phyllobacterium* was relatively enriched in glutathione transport genes and had relatively fewer genes related to secretion systems. The genus *Sphingomonas,* the second most abundant genera in *Madracis*, had one representative bMAG encoding prophages, Type IV secretion system genes, antibiotic resistance genes, detoxification genes, and a variety of phage defense mechanisms. The importance of secretion systems and pilus in these bMAGs is consistent with the overall functional gene analysis: among the 15 most differentially abundant COG categories were those related to secretion systems and pilus biosynthesis (type II secretion/type IV pili, type III and IV secretion systems, Flp/Tad pili), in addition to metabolism and energy production (non-phosphorylated Entner-Doudoroff pathway, NADH dehydrogenase), biosynthesis of essential molecules (isoleucine, leucine, and valine biosynthesis; aromatic amino acid biosynthesis; lysine, serine, purine, cobalamin (B12), molybdopterin, and phospholipid biosynthesis), and protein synthesis (50S ribosomal subunit) (Fig. S8A). Biosynthetic pathways were enriched in *Madracis*, while secretion systems and host attachment-related genes were more abundant in other corals.

All identified KEGG metabolism pathways were present in both *Madracis* and other corals. Among the 15 most differentially abundant metabolism-related KEGG pathways, 14 were more abundant in *Madracis* than in other corals (Fig. S8B). These pathways were involved in carbohydrate metabolism (e.g., fructose and mannose metabolism [ko00051] and glyoxylate and dicarboxylate metabolism [ko00630]), lipid and membrane metabolism (e.g., phospholipid biosynthesis [ko00564] and sphingolipid metabolism [ko00600]), amino acid and nitrogen metabolism (e.g., isoleucine, leucine, and valine biosynthesis [ko00290] and cysteine and methionine metabolism [ko00270]), cofactor and vitamin metabolism (e.g., cobalamin (B12) biosynthesis [ko00520] and riboflavin metabolism [ko00740]), and energy & redox metabolism (e.g., methane metabolism [ko00680], NADH dehydrogenase [ko00194], and sulfur metabolism [ko00920]). The only exception was riboflavin metabolism [ko00740], which was more abundant in other corals.

### Viral communities associated with Madracis

Viral communities were significantly different between *Madracis*, other coral species, and the CBL (F(2, 61) = 10.126, p = 0.001; Fig. 5A). Beta dispersion between groups only differed between other coral species and the CBL (p = 0.01, Tukey HSD) (Fig. S6). The Shannon diversity of viral communities in *Madracis* (5.53 ± 0.17) and the CBL (5.18 ± 0.13) were higher than that of other coral species (4.62 ± 0.18; Fig. 5B; Table S7), while the Simpson diversity of viral communities was similar between *Madracis* (0.99 ± 0.003) and the CBL (0.99 ± 0.002) or other coral species (0.93 ± 0.02). Viral community richness differed among groups (F(2, 61) = 58.85, p = 5.79e-15) and was 1.6- to 3.8-fold higher in *Madracis* (1500.25 ± 71.91) than in other corals (934.45 ± 56.42) and the CBL (397.95 ± 56.03). The evenness of the viral communities was similar between the two coral groups (0.76 ± 0.02 for *Madracis* and 0.69 ± 0.03 for other coral species) and lower than the CBL (0.90 ± 0.01; F(2, 61) = 19.98, p = 2.12e-07).

**Figure 5.**
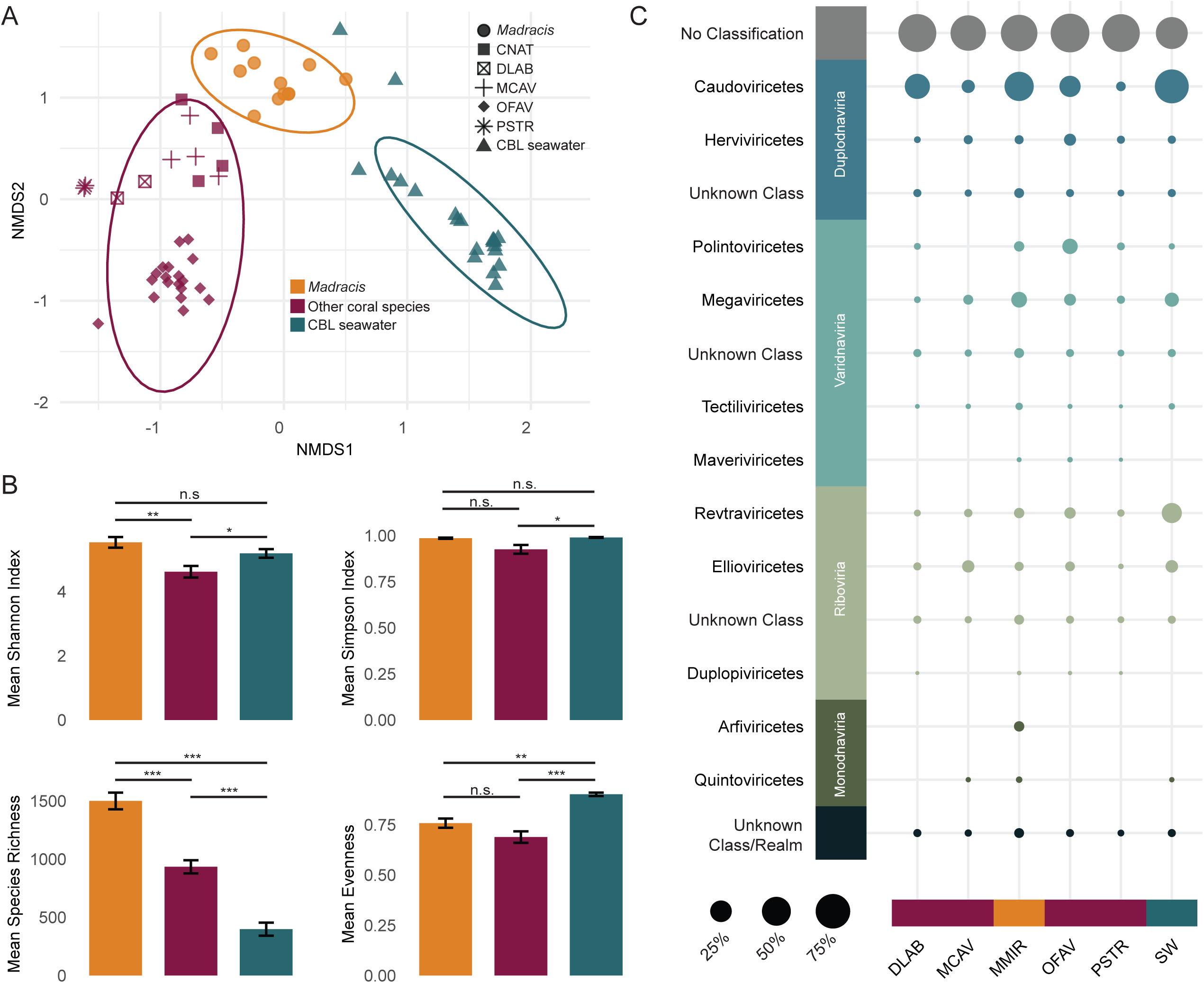
Viral community distinctions between *Madracis*, all other coral species in this study, and coral boundary layer (CBL) seawater. (A) Non-metric multidimensional scaling (NMDS) plot by viruses. NMDS plot is based on Bray Curtis distances calculated from the relative abundance of reads and utilizes 999 permutations. Circles denote a 95% CI for the clustered points. (B) Mean diversity indices of viral communities for each of the three groups, including Shannon Index, Simpson Index, Evenness, and Richness of bacterial genera. Significance codes: 0 ‘***’, 0.001 ‘**’, 0.01 ‘*’. (C) Bubble plot displaying the normalized relative fractional abundances of all identified viral classes by coral species or coral boundary layer (CBL). The size of the circles indicates the percent relative abundance of each class, and the color of the bubbles indicates viral realms. Coral species names are abbreviated as follows: *Colpophyllia natans* (CNAT), *Diploria labyrinthiformis* (DLAB), *Montastraea cavernosa* (MCAV), *Madracis aurentenra* (MAUR), *Orbicella faveolata* (OFAV), *Meandrina meandrites* (MMEA), and *Pseudodiploria strigosa* (PSTR).

A total of 2,820 dereplicated and quality-filtered viral genomes and genome fragments spanning 4 viral realms and 11 viral classes were identified in coral and CBL communities (Fig. 5C). The viral class *Caudoviricetes* dominated in *Madracis*, comprising 24.56% of the community, followed by *Megaviricetes* (3.33%) and *Retraviricetes* (1.59%). In other coral species, *Caudoviricetes* viruses were also the most abundant class, though at a 2.7-fold lower relative abundance (9.09%). These were more closely followed by viruses from the classes *Polintoviricetes* (5.55%), *Herviviricetes* (2.78%), and *Revtraviricetes* (2.56%). Of these viruses, 1,193 significantly contributed to the between-group differences, 179 of which explained 75% of the total dissimilarity. The 20 viruses contributing most to between-group differences, accounting for 22% of total dissimilarity, included viruses from the classes *Caudoviricetes*, *Megaviricetes*, and *Polintoviricetes*, as well as 9 viruses that could not be taxonomically classified (Table S14).

Proviruses accounted for an average of 6.75% ± 0.72% of all viruses in *Madracis*, significantly more than the 2.51% ± 0.41% of viruses in other coral species and 0.46% ± 0.11% of viruses in the CBL (Fig. 6A; Table S8). Among the 1,193 viruses that significantly contributed to the between-group differences, proviruses accounted for an average of 5.84% ± 0.66% of viruses in *Madracis*, which was significantly greater than in other coral species (2.62% ± 0.69%) and the CBL (0.21% ± 0.11%) (Fig. 6B; Table S8). Proviruses from the two coral groups encoded genes classified as COG categories “Mobilome: prophages and transposons”, “Replication, recombination, and repair”, and “Defense mechanisms” (Fig. 6C). Lower abundance metabolism-related functional categories including “Nucleotide transport and metabolism”, “Coenzyme transport and metabolism”, and “Amino acid transport and metabolism” were enriched in *Madracis* proviruses relative to those of other coral. The categories “Cell wall/surface interactions”, “Posttranslational modification, protein turnover, chaperones”, and “translation” were enriched in the proviruses of other corals.

**Figure 6.**
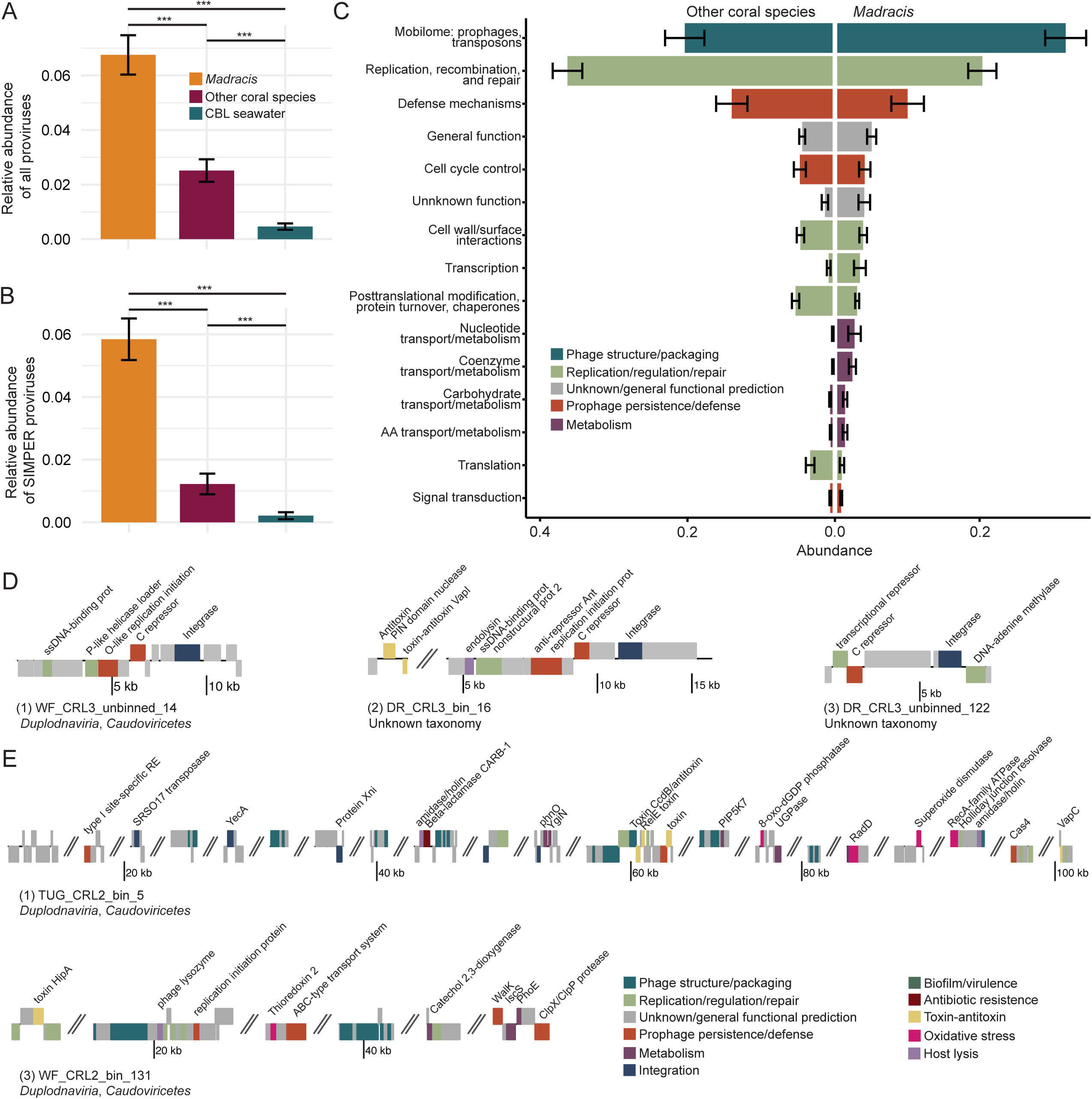
Genomic content of proviruses and viral indicators of the *Madracis* holobiont. (A) Relative abundance of proviruses among all viruses. (B) Relative abundance of proviruses among the SIMPER viruses. Significance codes: 0 ‘***’, 0.001 ‘**’, 0.01 ‘*’. (C) Top 15 Clusters of Orthologous Genes (COG) database functional categories of genes encoded by proviruses in other coral species (left) and *Madracis* (right). (D) Genome plots of *Madracis* indicator viruses encoding integrases. (E) Genome plots of two high-quality *Madracis* proviruses carrying auxiliary genes.

Forty-four viruses were significantly associated with *Madracis* at ≥ 95% specificity and fidelity (Table S15). High specificity values (e.g., A ≥ 0.95) represent viruses that are primarily or exclusively found in *Madracis* samples, and high fidelity values (e.g., B ≥ 0.95) represent viruses that are found across most or all *Madracis* samples. Of these 44 viruses, 10 with 100% specificity and fidelity in *Madracis* were selected for further genome analysis. Three encoded integrases, indicating that they may have the ability to integrate into their host genome, although they were not identified integrated in a bMAG (Fig. 6D). These temperate indicator viruses of *Madracis* primarily encode genes for genome replication and capsid structure. DR_CRL3_bin_16, the largest virus of the three, was an exception, encoding genes related to toxin-antitoxin systems. While only one of these viruses could be taxonomically classified (WF_CRL3_unbinned_14; *Caudoviricetes*), all three viruses also carry a phage repressor protein C which contains Cro/C1-type HTH and peptisase s24 domains. This regulator of the lysogenic-lytic switch is characteristic of lambda phage, a temperate member of the *Caudoviricetes* (67–69). In addition to these viruses, two high-quality prophage genomes encoded auxiliary gene products for antibiotic resistance (*CARB-1* beta-lactamase), metabolism (*phnO*, *ygiN,* Catechol 2,3-dioxygenase, *iscS*, *phoE*), prophage and host persistence (toxin-antitoxin systems, *recA*), and resilience under oxidative stress (8-oxo-dGDP phosphatase, *radD*, superoxide dismutase, thioredoxin 2) (Fig. 6E).

## DISCUSSION

### Coexistence with high microbial densities

In Curaçao, *Madracis mirabilis* forms expansive colonies composed of many tightly spaced branches, expanding on degraded and increasingly microbialized reefs where other corals have not achieved similar success (Fig 1). Microscopy results revealed significantly denser viral and microbial communities and increased virus-to-microbe ratios (VMRs) within *Madracis* branches (IB) compared to the boundary layer (BL) samples (Fig. 2). The natural branching complexity of *Madracis* forms a stagnant region in the center of the colony at high water flow (70), likely contributing to the increased densities of bacteria and viruses observed within the branches. The natural structure of *Madracis*, which harbors areas of dense microbial biomass, may have contributed to *Madracis*’ natural resistance to reef microbialization that is commonly associated with pollution and reef degradation, as observed in the Curacao reefs where *Madracis* occurs (9). This natural resistance to high bacterial densities may contribute to *Madracis* persistence and success in Curaçao.

### Disease resistance mechanisms

*Madracis* bacterial communities were distinct from those of other corals, showing dominance by *Ruegeria* and *Sphingomonas*, whereas other coral species were enriched in *Klebsiella*, Candidatus *Thioglobus*, and *Pseudomonas* (Fig. 3C). SIMPER analysis further identified *Ruegeria* and *Wenxinia* (family *Rhodobacteraceae*) as key genera differentiating *Madracis* from other coral species, with a relative depletion of *Nitrosospira* and *Beijerinckia* (Table S12). We identified four distinct *Ruegeria* bMAGs, three of which were enriched in Type VI secretion system (T6SS) genes and effectors that may contribute to *Madracis* success (Fig. 4). These genes are involved in interbacterial competition that may suppress invading pathogens, contributing to disease resistance in *Madracis*. Hcp proteins found in these bMAGs can function as effector toxins, targeting competing bacteria and promoting microbial dominance of beneficial taxa (71). Certain *Ruegeria* strains have been shown to inhibit coral pathogens like *Vibrio coralliilyticus* through unknown mechanisms (72, 73). Future studies should test in laboratory conditions if *Ruegeria*’s Hcp proteins can suppress *Vibrio coralliilyticus* and promote coral health.

*Madracis* also exhibited higher viral diversity and richness than in all other corals. *Caudoviricetes* (tailed phages) were the most abundant viral class across all corals, consistent with previous studies of coral viral communities, and were significantly more abundant in *Madracis* than in other corals (41, 74) (Fig. 5C). Tailed phages play roles in regulating bacterial populations, nutrient cycling, and genetic exchange (35). The elevated presence of these bacteria-infecting viruses in *Madracis* indicates that this species’ viral community may be more prominent in modulating bacterial community dynamics than in other corals. These bacteriophages could help control pathogen invasion via lytic infections, aligning with the Bacteriophage Adherence to Mucus (BAM) model (34).

*Madracis* microbiomes harbored a significantly higher abundance of prophages compared to other coral species (Fig. 6A; Fig. 6B). Genes related to toxin-antitoxin (TA) systems were prevalent in prophage genomes, highlighting a potential mechanism for *Madracis* microbiomes to resist bacteriophage superinfection and enhance bacterial stability under stressful conditions (75, 76). This aligns with previous work suggesting that TA systems enable bacterial “persisters” to survive harsh conditions, such as antibiotic exposure or nutrient depletion, by temporarily shutting down key processes (e.g., bacteriophage propagation) (77). Prophages infecting beneficial bacteria, such as *Ruegeria*, could help stabilize the bacterial community by preventing competitive displacement and increasing host resilience. Additionally, prophages carrying antibiotic-resistance genes, such as *CARB-1 beta-lactamase*, may confer protection against antibiotics and antimicrobial compounds produced by competitors, further enhancing bacterial persistence within the coral holobiont (78). Notably, three of the four *Ruegeria* bMAGs contained integrated proviruses, further supporting their role in microbiome resilience.

### Bacterial adaptations to nitrogen loading, organic matter inputs, and pollutants

Consistent with prior observations, coral-associated bacterial communities exhibited greater Shannon and Simpson diversity compared to seawater collected around the corals, underscoring the distinctions between these free-living and host-associated communities (79, 80) (Fig. 3B). Among corals, *Madracis* harbored a significantly richer bacterial community at the genus level than all other coral species combined. Microbiome diversity has been implicated in host responses to changing environments (81, 82). In the context of the success of *Madracis* in Curaçao, these highly diverse communities could provide advantages such as functional redundancy, metabolic versatility, and resilience in the face of environmental stress (83–85).

The *Ruegeria* genomes encoded genes related to nutrient cycling, organic matter degradation, and interbacterial competition. Consistent with other *Ruegeria* (86, 87), these genomes also contained genes associated with nitrogen fixation and metabolism (e.g., *fixH*, *ntrB*, *glnB*/*glnK*), as well as organic matter and pollutant degradation (e.g., *ssuD*, cytochrome P450, catechol 2,3-dioxygenase (C23O)). The presence of these genes suggests that *Ruegeria* may play a role in the bioavailability of nutrients for the coral host and its algal symbionts while also impacting organic matter availability for copiotrophic bacteria.

Another genus found to be highly abundant in *Madracis* was *Sphingomonas* (class *Alphaproteobacteria*) which has recognized roles in degradation of complex organic compounds, production of UV-protective carotenoids, and bioremediation (88–90) (Fig. 4). In degraded reef environments, microbial communities must adapt to increased organic pollution, oxidative stress, and shifting nutrient dynamics. A single representative *Sphingomonas* bMAG encoded Type IV secretion systems (T4SS), antibiotic resistance genes (e.g. *ampG*, β-lactamases), and multiple genes linked to bioremediation (e.g., catechol 2,3-dioxygenase, ring-hydroxylating dioxygenases, and arsenical resistance genes). This suggests that *Sphingomonas* contributes to detoxification processes and holobiont resilience in degraded reef environments, although its presence is not unique to *Madracis*. T4SS can facilitate horizontal gene transfer, potentially spreading antibiotic resistance, and other contact-based interactions, indicating broader ecological functions (91).

### Prophage contributions to bacterial persistence and metabolic diversity

As agents of gene transfer, proviruses contribute to the genomic flexibility, adaptability, and resilience of bacterial communities (35). Rather than causing lysis of bacterial populations, temperate viruses can integrate as dormant prophages, which may help maintain a stable and beneficial microbiome by introducing functional genes that enhance bacterial survival under environmental stress (92). If these prophages infect key symbionts, they may contribute to overall microbiome stability by maintaining beneficial bacterial populations in stressful reef environments. The enrichment of prophages in *Madracis* suggests that viral-mediated gene transfer may influence bacterial functions with direct implications for the coral host, particularly in degraded reef environments. Relative to other corals, *Madracis*-associated prophages encoded more genes related to metabolic functions (Fig. 6C), suggesting they may play roles in diversifying the metabolic potential of *Madracis*-associated bacteria.

*Madracis*-associated prophages genes may enhance bacterial survival in nutrient-rich, degraded environments. These genes included *phnO* which is involved in the metabolism of alternative phosphorus sources and could be beneficial in situations where nitrogen loading results in phosphorus limitation (93, 94). Additionally, genes encoding for enzymes like *ygiN*, which help detoxify quinone-like compounds, and catechol 2,3-dioxygenase, which facilitates the breakdown of aromatic compounds, can further contribute to the host’s metabolic flexibility (95, 96). Prophages encoding cysteine desulfurase enzymes may also be involved in sulfur metabolism and oxidative stress responses (97). Genes related to ROS detoxification (8-oxo-dGDP phosphatase*, radD,* superoxide dismutase (SOD), and thioredoxin 2 (*trx2*)) likely help maintain bacterial function under oxidative stress (98–101). Prophage genomes encoding these genes were enriched in *Madracis* relative to other coral species. Alternatively, these genes may play a role in protecting intracellular bacteria from oxidative stress caused by Symbiodiniaceae photosynthesis (102).

## Conclusion

Our study highlights the unique microbial traits of the *Madracis mirabilis* holobiont that may contribute to its success in degraded reef environments where other coral species struggle. The dense microbial and viral communities within *Madracis* colonies suggest a complex interplay between the coral and its microbiome, which may provide resilience against microbialization. The dominance of specific microbial genera, such as *Ruegeria*, and the presence of functional genes related to pathogen suppression, nutrient cycling, and pollutant degradation, indicate that *Madracis* may rely on its microbiome to support its survival in eutrophic conditions. Additionally, the high abundance of prophages and high viral diversity in *Madracis* suggest that viral-mediated gene transfer plays a role in maintaining a stable and adaptive microbiome. These findings underscore the potential of *Madracis* to serve as a model for understanding coral resilience in the face of anthropogenic reef degradation.

## ACKNOWLEDGEMENTS

We would like to thank Lars ter Horst, Maya Powell, and the Carmabi Foundation for logistical support and assistance in the field.

## AUTHOR CONTRIBUTIONS

CBS conceptualized the study. BAW, NSV, and AKS conducted field sampling and sample processing. MJAV selected field sites, facilitated sampling, and interpreted results. BAW conducted data retrieval, pipeline development, and analyses. BAW wrote the first draft, and all authors edited the manuscript.

## Footnotes

1 We acknowledge the proposed renaming of *Madracis mirabilis* to *Madracis auretenra*. However, due to ongoing concerns over the validity of the type specimen used in the revision by Locke et al (2007) and the widespread and longstanding use of *M. mirabilis* in the literature, we continue to use *M. mirabilis* in this manuscript for clarity and consistency.

